# The evolving systemic biomarker milieu in obese ZSF1 rat model of human cardiometabolic syndrome: Characterization of the model and cardioprotective effect of GDF15

**DOI:** 10.1101/2020.03.20.000307

**Authors:** Marina Stolina, Xin Luo, Denise Dwyer, Chun-Ya Han, Rhonda Chen, Ying Zhang, YuMei Xiong, Yinhong Chen, Jun Yin, Brandon Ason, Clarence Hale, Murielle M. Véniant

## Abstract

Cardiometabolic syndrome has become a global health issue. Heart failure is a common comorbidity of cardiometabolic syndrome. Successful drug development to prevent cardiometabolic syndrome and associated comorbidities requires preclinical models predictive of human conditions. To characterize the heart failure component of cardiometabolic syndrome, cardiometabolic, metabolic, and renal biomarkers were evaluated in obese and lean ZSF1 20-to 22-week-old male rats. Cardiac function, exercise capacity, and left ventricular gene expression were also analyzed. Obese ZSF1 rats exhibited multiple features of human cardiometabolic syndrome by pathological changes in systemic renal, metabolic, and cardiovascular disease circulating biomarkers. Hemodynamic assessment, echocardiography, and decreased exercise capacity confirmed heart failure with preserved ejection fraction. RNA-seq results demonstrated changes in left ventricular gene expression associated with fatty acid and branched chain amino acid metabolism, cardiomyopathy, cardiac hypertrophy, and heart failure. Twelve weeks of growth differentiation factor 15 (GDF15) treatment significantly decreased body weight, food intake, blood glucose, and triglycerides and improved exercise capacity in obese ZSF1 males. Systemic cardiovascular injury markers were significantly lower in GDF15-treated obese ZSF1 rats. Obese ZSF1 male rats represent a preclinical model for human cardiometabolic syndrome with established heart failure with preserved ejection fraction. GDF15 treatment mediated dietary response and demonstrated a cardioprotective effect in obese ZSF1 rats.

## Introduction

Cardiometabolic syndrome (CMS)—a condition that encompasses impaired metabolism (insulin resistance [IR], impaired glucose tolerance), dyslipidemia, hypertension, renal dysfunction, central obesity, and heart failure (HF)—is now recognized as a disease by the World Health Organization (WHO) and the American Society of Endocrinology [1]. Obesity and diabetes mellitus comorbidities are associated with progressive left ventricular (LV) remodeling and dysfunction. Also, these comorbidities are commonly observed in HF with preserved ejection fraction (HFpEF) [2]. Results from a recent epidemiological study (cohort of 3.5 million individuals) demonstrated an incremental increase in the hazard ratio (HR) for HF; HRs were 1.8 in normal weight individuals with three metabolic abnormalities, 2.1 in overweight individuals with three metabolic abnormalities, and up to 3.9 in obese individuals with three metabolic abnormalities. Incidence of HFpEF, which currently represents approximately 50% of all HF cases, continues to rise and its prognosis fails to improve partly due to the lack of therapies available to treat this disease [3].

An important step in the development of novel therapeutic agents against CMS is the establishment of a preclinical model that represents a cluster of cardiometabolic disturbances that are similar to those of the human condition. The obese ZSF1 rat model (generated by crossing lean, non-hypertensive, female Zucker diabetic fatty rats [ZDF, +/*fa*] with lean, spontaneously hypertensive, HF-prone male rats [SHHF/Mcc, +/*facp*])[4] exhibits features and complications that resemble what is observed in human CMS [5, 6]. Twenty-week-old obese ZSF1 male rats developed diastolic dysfunction based on prolonged τ and elevated end-diastolic pressure-volume relationship (EDPVR), and showed exercise intolerance, which is an important feature of human HFpEF [7]. Although both lean and obese ZSF1 rats, by inheritance of a hypertensive gene, showed elevated blood pressure [4], only 20-week-old obese ZSF1 males demonstrated LV hypertrophy, left atrial (LA) dilation, and increased myocardial stiffness due to myofilament changes [8].

Growth differentiation factor 15 (GDF15), also called macrophage inhibitory cytokine (MIC-1), is a distant member of the transforming growth factor β (TGF-β) superfamily. GDF15 is a homodimeric secreted protein with a mass of 25 kDa [9, 10]. Circulating levels are increased in humans with metabolic syndrome [11] and in those with increased risk of cardiovascular disease (CVD) [12, 13]. Recently published studies in obese preclinical models demonstrated an aversive dietary response to GDF15 treatment, leading to an improvement in metabolic parameters. Thus, daily injections of GDF15 to mice for 14 days to 21 weeks resulted in significantly reduced body weight and food intake, increased energy expenditure, improved glucose tolerance, and reduced inflammatory cytokines [14]. Administration of human GDF15 to rodents via an adenovirus system and to obese monkeys via protein injections led to body weight loss and an improved metabolic profile [15]. Treatment of obese mice with a human GDF15-Fc fusion protein (Fc-GDF15) led to reduced appetite and body weight and a shift of metabolic parameters toward lipid oxidation [15–18]. Weekly administration of Fc-GDF15 to obese cynomolgus monkeys for 28–42 days resulted in significantly reduced body weight and food consumption, lower serum triglyceride levels, and improved serum insulin [15, 19]. To date, GDF15 is shown to have mediated aversive dietary response, influenced the governance of systemic energy balance, and prevented obesity through enhanced thermogenesis and oxidative metabolism [20, 21].

While impaired metabolism, LV hypertrophy, LA dilation, increased myocardial stiffness, and decreased exercise capacity of obese ZSF1 males have been published in several separate scientific reports [5, 8, 22], the obesity-induced changes in systemic cardiovascular protein biomarkers and LV gene expression have not been reported. In the current study, we performed a comprehensive comparative evaluation of 20-to 22-week-old male lean and obese ZSF1 rats by systematically studying the metabolic, renal, and cardiovascular protein biomarkers. We also used echocardiography, invasive hemodynamic assessment, exercise capacity assessment, and LV gene expression analysis to complete the characterization of this animal model. Additionally, we investigated whether 12 weeks of treatment with Fc-GDF15 in 22-week-old obese ZSF1 males would result in decreased body weight and food intake and improvement in metabolic profile, similar to prior non-clinical studies [15, 18], and also whether a cardioprotective effect could be demonstrated by improving exercise capacity and circulating levels of cardiovascular protein biomarkers. Overall, our study suggests that obese ZSF1 male rats represent a preclinical model for human cardiometabolic syndrome with established HFpEF.

## Materials and methods

### Animal welfare and husbandry

All studies were performed in accordance with the Institutional Animal Care and Use Committee guidelines and complied with the Final Rules of the Animal Welfare Act regulations (Code of Federal Regulations, Title 9), the Public Health Service Policy on Humane Care and Use of Laboratory Animals in the Office of Laboratory Animal Welfare (2002), and the Guide for the Care and Use of Laboratory Animals from the National Research Council (1996). All rodent studies were conducted at Amgen Inc and were approved by the Amgen Institutional Animal Care and Use Committee (IACUC). Animals were maintained in rooms with a 12-hour light/dark cycle, temperature of 22°C, and humidity of 30–70%. Animals had free access to food and water and were maintained on standard rodent chow unless otherwise indicated. Rats were single-housed at Amgen’s Association for Assessment and Accreditation of Laboratory Animal Care (AAALAC)–accredited facility in filter-top cages on corn cob bedding, with ad libitum access to pelleted feed (Harlan/Teklad Irradiated Global Soy Protein-Free Extruded Rodent Diet 2920x; Harlan, Madison, WI, USA) and reverse-osmosis purified water via an automatic watering system. Animals were maintained in pathogen-free conditions with a 12-hour light/dark cycle and had access to enrichment opportunities.

### In vivo study design for characterization of obese ZSF1 rat as a preclinical model of human CMS

Eighteen-week-old lean ZSF1 male (strain 379, *n* = 14) and obese ZSF1 (strain 378, *n* = 14) rats were purchased from Charles River Laboratories (Kingston, NY, USA) and single-housed at Amgen’s AAALAC-accredited facility. After 2 weeks of acclimation, both groups of rats (*n* = 8 per group) were subjected to blood collection via tail vein, echocardiography, hemodynamic assessment, and evaluation of exercise endurance (time and distance) using a treadmill. A separate cohort from the same batch of acclimated obese ZSF1 (*n* = 6) and lean ZSF1 (*n* = 6) rats was subjected to heart isolation, RNA extraction, and gene expression analysis.

### In vivo study design for the evaluation of Fc-GDF15 treatment effect on CMS-specific biomarkers in obese ZSF1 rats

Twenty-week-old obese ZSF1 male rats (strain 378, *n* = 30) were purchased from Charles River Laboratories, single-housed at Amgen’s AAALAC-accredited facility, and acclimated for 2 weeks. At 22 weeks of age, baseline blood collection (for metabolic biomarker evaluation) and body weight assessment were performed for every animal. Twenty-four hours later, the rats were randomized into two groups and injected subcutaneously once a week for 12 weeks with either A5.2Su buffer (vehicle group, *n* = 15) or 1.5 mg/kg DhCpmFc-(G4S)4-hGDF15 [15] (Fc-GDF15 group, *n* = 15). For the duration of the study, every rat was subjected to weekly blood collection, food intake, and body weight assessments. At the end of the study, rats from both treatment groups were subjected to echocardiography, hemodynamic assessment, and evaluation of exercise endurance (time and distance) using treadmill equipment.

### Cardiovascular, kidney injury, and metabolic biomarkers in serum/plasma

Animals were fasted for 4 hours prior to blood collection. The first 0.5 mL aliquot of whole blood was collected from the tail vein into serum separator tubes (Microtainer, Becton Dickinson, Franklin Lakes, NJ, USA), and the second 0.5 mL aliquot of whole blood into ethylenediaminetetraacetic acid (EDTA) plasma separation tubes (Microtainer, Becton Dickinson, Franklin Lakes, NJ). Separated serum/plasma was aliquoted and stored at –80°C. Blood glucose levels were measured using AlphaTrak 2 glucose strip (Abbott Laboratories, Lake Bluff, IL, USA). Blood insulin was evaluated using a rat insulin enzyme-linked immunosorbent assay (ELISA) kit (Alpco, Salem, NH, USA). Serum triglyceride and cholesterol levels were measured weekly by triglyceride quantitation kits (Fisher Diagnostics, Middletown, VA, USA). Systemic levels of GDF15 (R&D Systems, Minneapolis, MN, USA) and proinsulin (Mercodia, Uppsala, Sweden) in serum/plasma of 20-week-old ZSF1 male rats were evaluated by commercial ELISAs. A limited array of circulating metabolic hormones (amylin [active], C-peptide 2, ghrelin [active], gastric inhibitory polypeptide [GIP, total], glucagon-like peptide 1 [GLP-1, active], glucagon, interleukin-6 [IL-6], leptin, pancreatic polypeptide [PP], and peptide YY [PYY]) was assessed in serum/plasma collected from lean and obese ZSF1 rats at the age of 20 weeks by rat-specific MILLIplex kit (Millipore, Billerica, MA, USA). Kidney injury markers KIM-1 and NGAL were evaluated in serum from 20-week-old lean and obese ZSF1 rats by multiplex MILLIplex kits (Millipore). Serum NT-proBNP and rat cardiac injury markers (fatty-acid-binding protein 3 [FABP3] and myosin light chain 3 [Myl3]) were measured by using rat-specific single-plex or multiplex commercial assays from Meso Scale Diagnostics (Rockville, MD, USA). Systemic levels of vascular markers were evaluated at the end of the Fc-GDF15 and vehicle treatment of obese ZSF1 rats by using multiplex MILLIplex kits (Millipore). The rat vascular injury panels included VEGF, MCP1, TIMP1, TNFα, vWF, adiponectin, sE-selectin, and sICAM1. Systemic aldosterone levels were measured by enzyme immunoassay (m/r/h Aldosterone ELISA; Enzo Life Sciences, Farmingdale, NY, USA) and serum osteopontin (OPN) by Rat Single Plex Kit (Millipore). All the assays were performed in accordance with manufacturer protocols.

### RNA isolation and sequencing analysis of left heart

At scheduled necropsy, LA and LV of the heart were washed briefly in RNase-free saline, placed in cryo-tubes, snap-frozen in liquid nitrogen, and stored at –80°C. Samples were subjected to dry pulverization before being homogenized in buffer (350 μL of Qiagen RLT buffer with 1% β-mercaptoethanol; Qiagen, Germantown, MD, USA). The homogenate was transferred to an RNase-free 1.5 mL centrifuge tube, after which 590 μL RNase-free water and 10 μL of a 20 mg/mL proteinase K solution were added. Samples were incubated at 55°C for 10 minutes, centrifuged, and the supernatant collected. RNA was then extracted using RNeasy Micro Kit (Qiagen) with on-column DNase treatment (Qiagen) according to the manufacturer’s instructions. RNA concentration and integrity were assessed using a Bioanalyzer (Agilent, Santa Clara, CA, USA). Samples with ≥80 ng total RNA and RNA integrity numbers (RIN) ≥7 were used for sequencing. Raw reads were processed using OmicSoft (Cary, NC, USA) Array Studio software (Oshell.exe v10.0) [23]. The expression level was expressed as fragments per kilobase per million quantile normalized (FPKQ) values, which were generated using OmicSoft’s implementation of the RSEM algorithm and normalized using upper-quartile normalization [24].

### Differential expression and pathway analysis

Differential expression analysis was performed using DESeq2 v1.10.1 for obese vs lean ZSF1 LV comparison [25], and gene expression fold changes were calculated using FPKQ values. Genes with Benjamini-Hochberg corrected *p*-value < 0.05 and fold change between the two conditions ≥ 1.5 or ≤ 0.67 were selected as significantly differentially expressed genes (DEGs).

The analysis of the genes that have human homologue and are abundant in human heart (top three out of 45 tissue roll-ups with median FPKQ *≥* 1) was based on Genotype-Tissue Expression (GTEX) data [26]. Heart abundant altered genes were visualized in a graphic heatmap using the ComplexHeatmap R package [27]. All the DEGs and heart abundant DEGs were further annotated by ingenuity pathway analysis (IPA; QIAGEN, Redwood City, CA, USA). Metabolic signaling pathway enrichment was done on the significantly DEGs by an adjusted *p*-value cutoff of 0.05 by IPA. Gene expression enrichment in diseases was analyzed by using Medical Subject Headings (MeSH) terms using the meshes R package [28]. Dotplots and heatplots were generated with clusterProfiler [29].

### Echocardiography

Non-invasive echocardiograms were obtained on anesthetized rats (isoflurane, 3% induction, 1.5% maintenance) using a Vevo 2100 imaging system (FUJIFILM VisualSonics, Inc, Toronto, Canada). Animals were shaved and placed on a platform, and a thermo-couple probe was used to assess body temperature and to adjust the temperature of the platform to maintain normothermia. Sonography gel was applied on the thorax. Two-dimensional targeted B-mode and M-mode imaging was obtained from the long-axis and short-axis views at the level of the papillary muscle. Coronary flow was measured by pulsed-wave (PW) Doppler, and movement of the LV wall was measured by tissue Doppler by placing the probe to the chest and focusing the image on the ventricular wall region to achieve an apical four-chamber view. Animals were subsequently wiped clean of sonography gel, allowed to achieve consciousness, and returned to their home cage.

### Invasive hemodynamic assessment

Animals underwent general anesthesia using ketamine/diazepam (intraperitoneal) injection (80–100 mg/kg ketamine; 5–10 mg/kg diazepam) followed by a ketamine boost (10– 20 mg/kg) as necessary to maintain the anesthesia plane. Anesthetized animals had the surgical sites shaved with a small animal shear. Animals were then placed on a heated surgical platform, and body temperature was monitored and maintained throughout the study via a rectal probe. Arterial pressure was measured via femoral artery catheterization (SPR-839; Millar, Houston, TX, USA). An incision in the medial aspect of the leg was made, and the fat overlaying the femoral vessels and femoral nerve in the area between the abdominal wall and the upper leg was gently teased apart using forceps or Q-tips. The fascia overlying the artery, nerve, and vein was removed. The artery was isolated and ligated by a distal suture, and a loose tie was held in place using a hemostat. A microvessel clip or suture was placed on the artery near the abdominal wall, and a small incision was made into the vessel using Vannas micro-scissors or a 25-gauge needle. The clip was released by one hand, and the Millar catheter was advanced quickly into the femoral artery, followed by the tightening of the proximal suture to prevent blood loss. Pressure was then recorded. LV pressure and volume were measured via carotid artery catheterization (SPR-838; Millar). The right carotid artery was isolated, and a suture was tied around the vessel approximately 0.25 cm below the skull base to disrupt blood flow. A suture was placed around the artery near the rib cage and held in place using a hemostat. A small incision was made into the vessel using Vannas micro-scissors or a 25-gauge needle. The clip was released by one hand, and the Millar catheter was advanced quickly into the carotid artery, followed by the tightening of the proximal suture to prevent blood loss. The catheter was advanced until it entered the LV for pressure-volume loop measurement. Animals were euthanized under anesthesia after completion of the data acquisition.

### Treadmill assessment

Rats were subjected to evaluation of exercise endurance (time and distance) using a treadmill. Endurance exercise performance was estimated using two parameters: run duration (minutes) and distance (meters). Oxygen consumption, an indicator of exercise capacity, was measured by monitoring the O_2_ concentration of expired air. Peak chamber oxygen decrement, which usually occurs just prior to exhaustion, was used to calculate and define the animal’s peak volume of oxygen consumption (VO_2_). The contribution of anaerobic metabolism to overall energy production during exercise was estimated by calculating the respiratory exchange ratio (RER), the ratio of VO_2_ to VCO_2_ (the amount of CO_2_ within the chamber just prior to exhaustion). Rats were initially familiarized to the motorized rodent treadmill (Columbus Instruments, Columbus, OH, USA) for 5 days before completing the exercise performance test. Rats were placed on the treadmill and allowed to adapt to the surroundings for 2–5 minutes before starting. Familiarization runs consisted of 10 minutes of running on an incline of 10° at a speed of 12 m/min. For the treadmill challenge, rats were placed on the treadmill and allowed to adapt to the surroundings for 2–5 minutes before starting. The treadmill was initiated at a speed of 8.5 m/min with a 0° incline. After 3 minutes, the speed and incline were raised up to 10 m/min. The incline was subsequently increased progressively by 2.5 m/min every 3 minutes. The incline was progressively increased by 5° every 9 minutes to a maximum of up to 30°. Exercise continued until exhaustion, which is defined as the inability to maintain running speed despite repeated contact with the electric grid (three shocks in less than 15 seconds).

### Statistical analysis

Results for serum protein biomarkers, hemodynamic assessments, echocardiography, and systemic metabolic parameters are expressed as mean ± standard error of the mean (SEM). Differences between the two groups were examined between lean ZSF1 and obese ZSF1 rats of the same age or at the end of the Fc-GDF15 treatment (week 12) between vehicle-treated and Fc-GDF15–treated obese ZSF1 rats. Unpaired Student’s *t* test was performed to evaluate the statistical significance between the groups. A value of *p* < 0.05 was used to determine statistical significance. Stars (*) indicate significance: * *p* < 0.05, ** *p* < 0.01, *** *p* < 0.001, **** *p* < 0.0001. All the statistical analysis was performed by using GraphPad Prism (Version 7.04, GraphPad Software, San Diego, CA).

## Results

### At the age of 20 weeks, obese ZSF1 male rats exhibited increased body weight and impaired metabolism

We have confirmed previous reports [4] and demonstrated that 20-week-old obese ZSF1 rats exhibit impaired markers of metabolism (S1 Table; Fig 1), including elevated body weight (38% increase vs lean group; *p* < 0.0001) and significantly increased blood glucose (1.6-fold; *p* = 0.0060), serum insulin (14.7-fold; *p* = 0.0019), cholesterol (2.9-fold; *p* < 0.0001), and triglyceride (15-fold; *p* < 0.0001) levels relative to lean ZSF1 littermate controls. Significant decrease in serum adiponectin from 2.6 ± 0.09 μg/mL in lean ZSF1 serum to 1.9 ± 0.05 μg/mL in obese ZSF1 serum (*p* < 0.0001) indicated deposition of newly formed fat and served as a reliable obesity biomarker.

**Fig 1.**
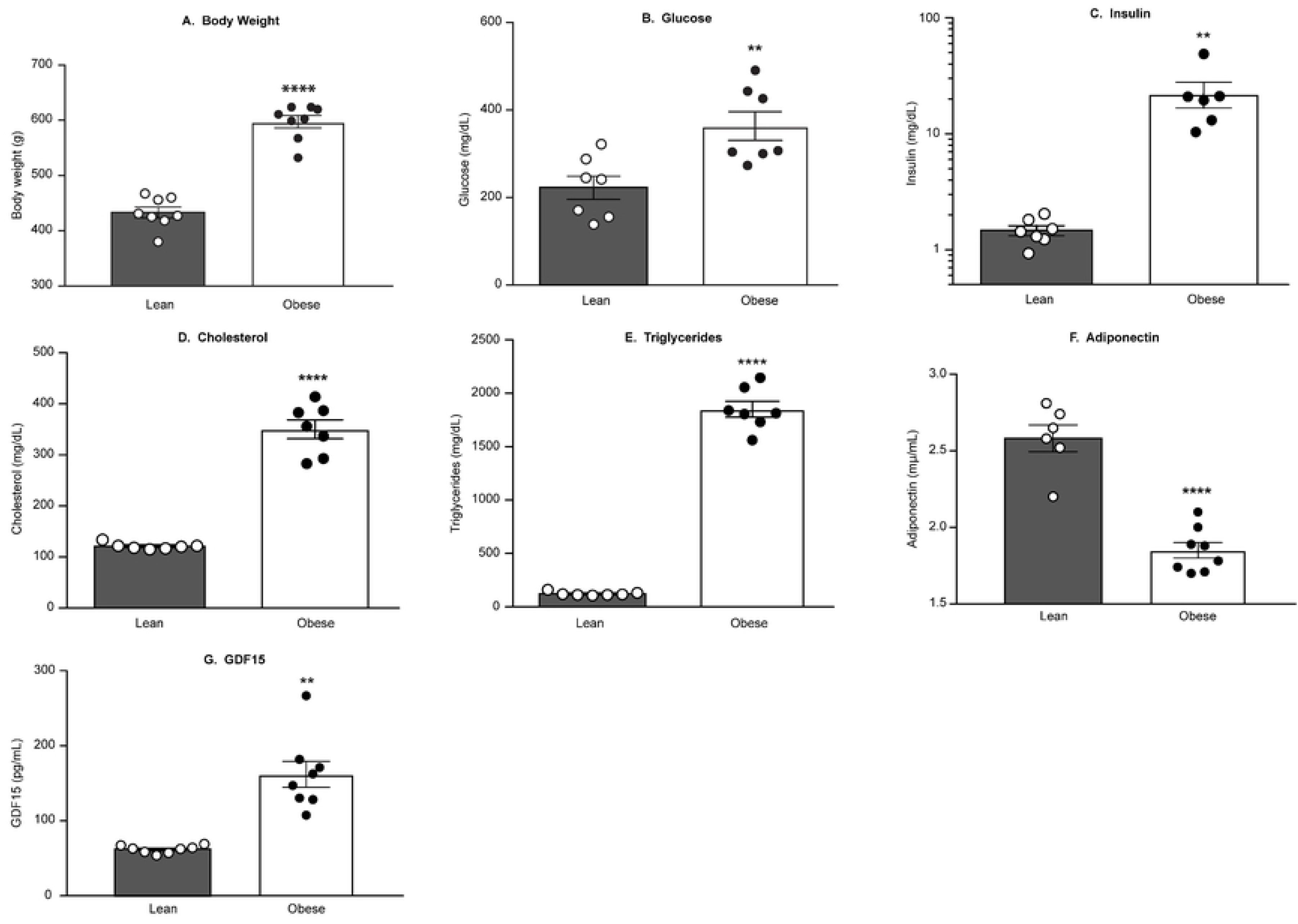
Obese ZSF1 male rats exhibited increased BW, glucose, insulin, cholesterol and triglycerides levels and decreased adiponectin and increased GDF15 levels. Obese ZSF1 male rats at the age of 20 weeks exhibited increased body weight (A), elevated blood glucose (B), insulin (C), cholesterol (D), and triglyceride levels (E), decreased adiponectin (F), and increased growth differentiation factor 15 (GDF15) (G). Stars (*) indicate significance (* *p* < 0.05, ** *p* < 0.01, *** *p* < 0.001, **** *p* < 0.0001) by unpaired two-tailed *t* test.

### Twenty-week-old obese ZSF1 male rats demonstrated pancreatic and renal dysfunction

Since the array of biomarker changes relevant to the progression and establishment of CMS in humans includes type 2 diabetes (T2D) and renal impairment, we evaluated systemic levels of major pancreatic and kidney injury biomarkers in obese ZSF1 rats compared with their lean littermates (S1 Table). Twenty-week-old obese ZSF1 male rats demonstrated pancreatic dysfunction by showing a 2.7-fold elevation in systemic C-peptide, a 47-fold increase in proinsulin (Fig 2B), a 6.3-fold increase in serum active amylin (Fig 2C), and elevated glucagon (37.8 ± 4.6 pg/mL in obese ZSF1 vs 25.8 ± 2.5 pg/mL in lean ZSF1 littermates; *p* = 0.0388; Fig 2D). Impaired renal function in the obese ZSF1 group was characterized with significantly increased serum kidney injury markers NGAL (1.5-fold vs lean ZSF1; Fig 2E), KIM-1 (2.2-fold vs lean ZSF1; Fig 2F), and clusterin (1.4-fold vs lean ZSF1; Fig 2G).

**Fig 2.**
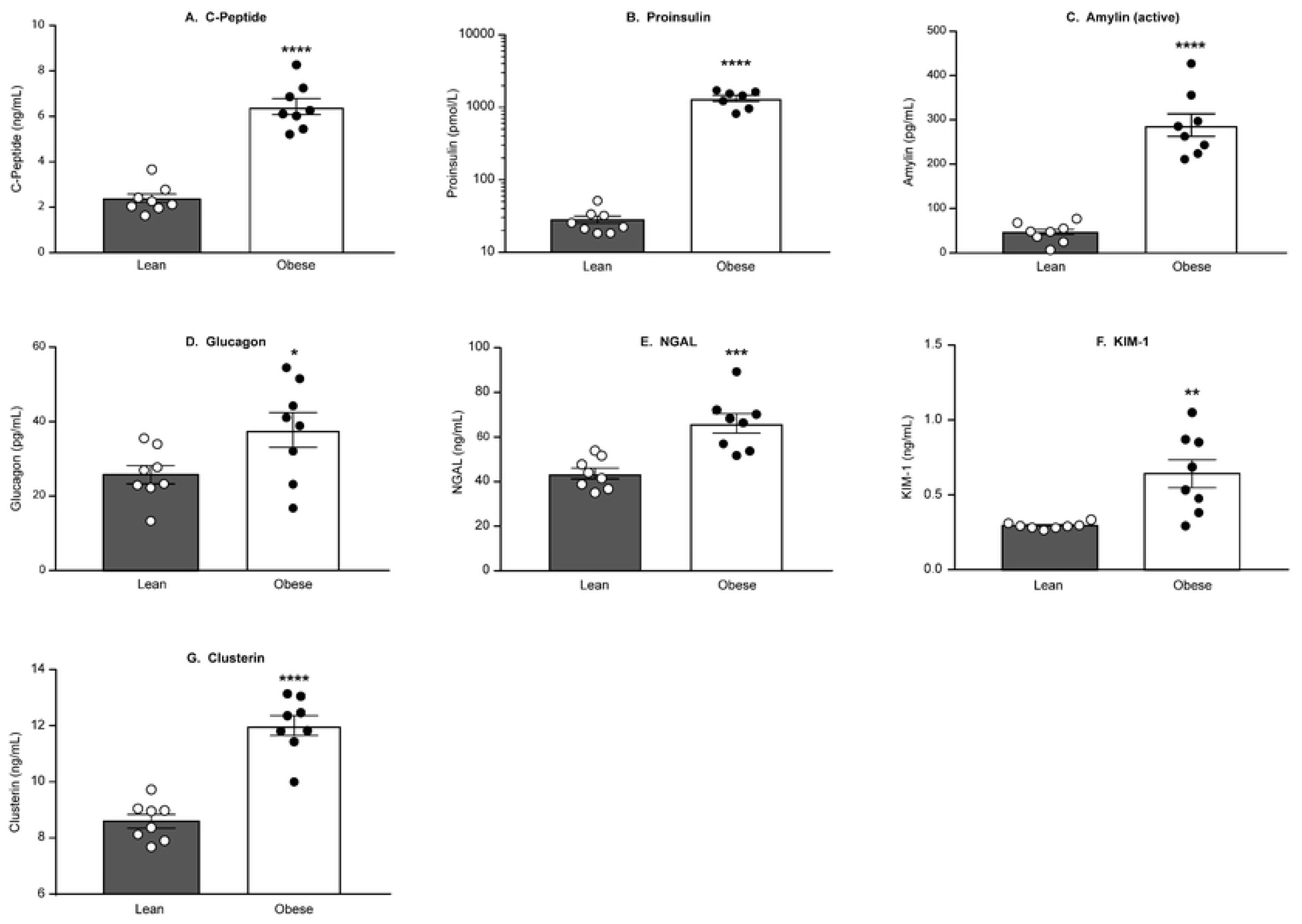
Obese ZSF1 rats demonstrated pancreatic dysfunction and impaired renal function. Obese ZSF1 rats show increased C-peptide (A), proinsulin (B), active amylin (C), and glucagon (D). Obese ZSF1 rats exhibit impaired renal function by showing increased serum levels of kidney injury markers NGAL (E), KIM-1 (F), and clusterin (G). Stars (*) indicate significance (* *p* < 0.05, ** *p* < 0.01, *** *p* < 0.001, **** *p* < 0.0001) by unpaired two-tailed *t* test.

### Obese ZSF1 male rats demonstrated impaired cardiac function and reduced exercise capacity at the age of 20–21 weeks

The complete set of invasive hemodynamic assessment, echocardiography, and exercise capacity of 20-to 21-week-old lean and obese ZSF1 male rats is presented in S3 Table. The heart-to-brain weight ratios were significantly higher in obese ZSF1 rats than in lean ZSF1 littermates (Fig 3A). Invasive hemodynamic assessment at 21 weeks for rats anesthetized with ketamine/diazepam showed an increased relaxation constant *tau* (Fig 3B). Echocardiography at 20 weeks under isoflurane anesthesia revealed that obese ZSF1 rats exhibit a significant decrease in heart rate (Fig 3C), ratio of mitral peak velocity of early filling to early diastolic mitral annular velocity (E/E’) (Fig 3D), and isovolumic relaxation time (IVRT) (Fig 3E), while maintaining a normal ejection fraction (Fig 3F). We further demonstrated that obese ZSF1 rats exhibit a limited exercise capacity with significantly shorter time to exhaustion (Fig 3H), lower peak VO_2_ (Fig 3I), and shorter distance (Fig 3G) following a treadmill exercise challenge.

**Fig 3.**
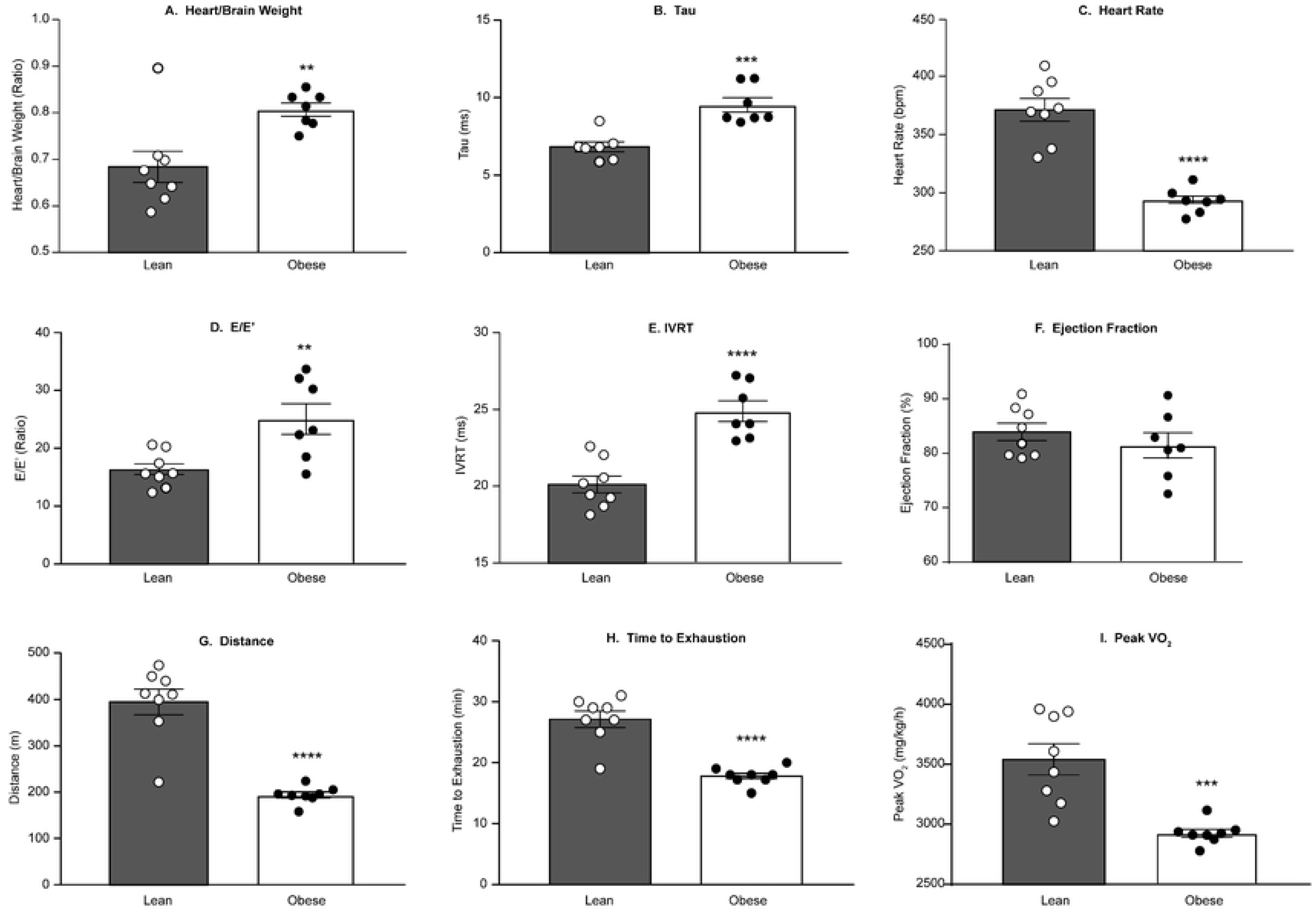
Obese ZSF1 rats exhibited diastolic dysfunction and decreased exercise capacity. The heart-to-brain weight ratio (by echocardiography) is increased in obese ZSF1 rats (A), and invasive hemodynamic assessment provides evidence of diastolic dysfunction by increased *tau* (B) when compared with lean ZSF1 rats. Echocardiography reveals that obese ZSF1 rats exhibit a decrease in heart rate (C) and diastolic dysfunction by increase in the E/E’ ratio (D) and isovolumetric relaxation time (IVRT) (E), while % ejection fraction is not different from lean ZSF1 (F). Obese ZSF1 rats exhibit decreased exercise capacity as measured by distance (G), time (H), and peak VO_2_ (I). Stars (*) indicate significance (* *p* < 0.05, ** *p* < 0.01, *** *p* < 0.001, **** *p* < 0.0001) by unpaired two-tailed *t* test.

### At 20 weeks of age, obese ZSF1 male rats exhibited significant changes in major systemic biomarkers of cardiovascular function

To evaluate the translational value of the obese ZSF1 rat model for human CMS, we next profiled major systemic CVD biomarkers (S2 Table; Fig 4A–4F) and compared the results with the gene expression of the same markers in the left heart of 20-week-old lean and obese ZSF1 male rats (Fig 4G and 4H). The array of systemic changes in CVD-related biomarkers included increased blood levels of hypertension marker aldosterone (Fig 4A), elevated FABP3, a biomarker of heart pathology (Fig 4B), and significantly increased vascular markers IL-16 (Fig 4C) and ST2 (Fig 4D). Systemic concentration of NT-proBNP (Fig 4E) in obese ZSF1 rats was significantly lower than in the lean rats, and OPN (Fig 4F) level in circulation was not different between lean and obese ZSF1 rats. Gene expression of the above markers in the left heart was not different between lean and obese ZSF1 rats for FABP3, IL-16, IL1RL1 (data not shown), and NPPB (NT-proBNP protein; Fig 4G), whereas the local SPP1 mRNA expression (OPN gene) in both LA and LV was significantly elevated in obese ZSF1 rats (Fig 4H).

**Fig 4.**
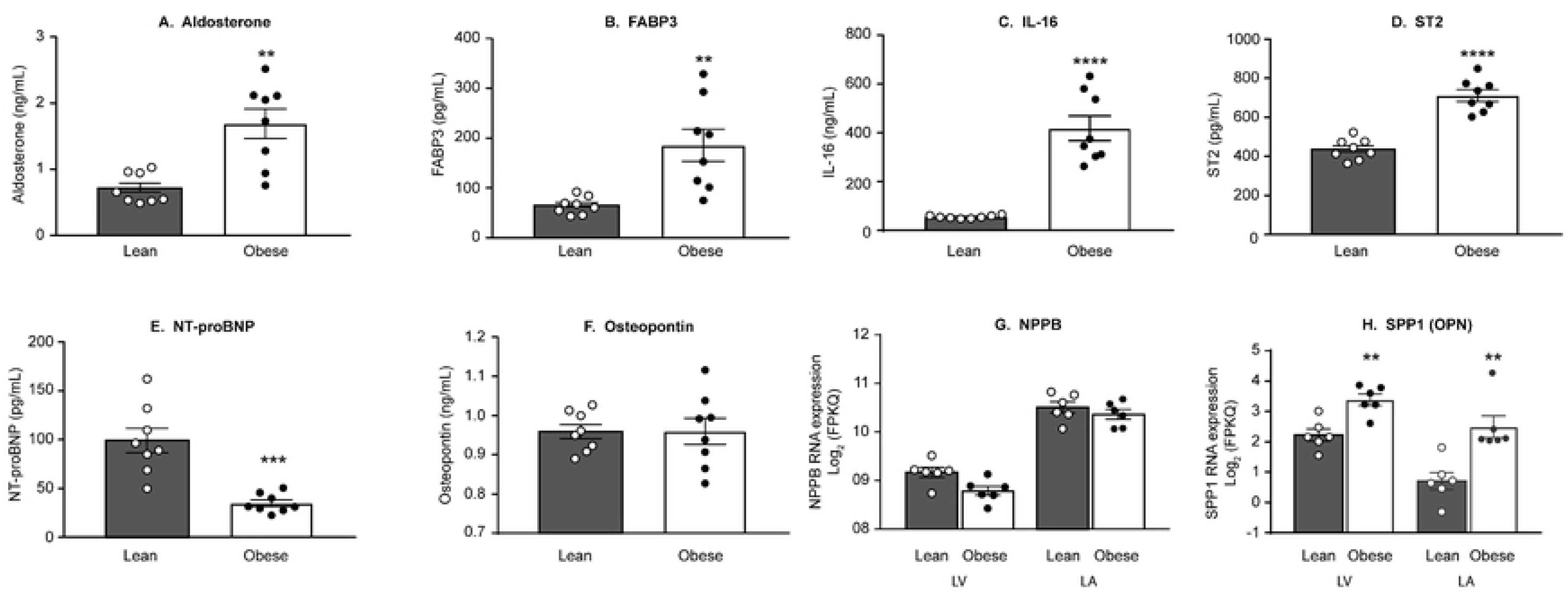
Twenty-week-old obese ZSF1 male rats exhibited cardiovascular dysfunction. **obese ZSF1** show increased blood levels of aldosterone (A), fatty-acid-binding protein 3 (FABP3; B), interleukin-16 (IL-16; C), and ST2 (D). In obese ZSF1 rats, systemic concentration of NT-proBNP (E) is lower compared with the lean cohort and osteopontin (OPN; F) level in circulation is not different between the two cohorts. The mRNA expression of NPPB (NT-proBNP protein; G) in left heart was not different between obese ZSF1 and lean ZSF1 groups, whereas the local SPP1 (OPN) expression in left atria (LA) and left ventricle (LV) (H) was significantly elevated in obese ZSF1 rats. Stars (*) indicate significance (* *p* < 0.05, ** *p* < 0.01, *** *p* < 0.001, **** *p* < 0.0001) by unpaired two-tailed *t* test.

### CMS-related gene expression changes in the left heart of 20-week-old male ZSF1 rats

To characterize the transcriptional effects of obesity and metabolic dysfunction on the expression of left heart specific/abundant genes, RNA-seq was performed on LV isolated from 20-week-old lean and obese ZSF1 male rats (Fig 5). Comparison of heart abundant protein coding gene expression levels in LV biopsies from lean and obese ZSF1 rats revealed a relatively small number of genotype-driven gene expression changes (Fig 5A): the expression of 56 genes was significantly increased (FC > 1.5; *p* < 0.05; S6 Table), and the expression of 48 genes was significantly decreased (FC < 0.75; *p* < 0.05; S7 Table) in obese rats vs lean rats. The IPA pathway enrichment analysis of significantly dysregulated genes in the LV of obese ZSF1 rats revealed that the altered genes are significantly enriched in fatty acid and branched-chain amino acid (BCAA) metabolism pathways (Fig 5B). The increase in ACADM, EHHADH, HADHA, and HADHB gene expression was shared by both fatty acid and BCAA metabolism pathways. Altered ACAA2, ACOT2, ACSL6, ECI1, BDH1, HMGCS2, and IDI1 gene expression was fatty acid metabolism specific, and altered expression of DUSP26, MAOA, MAOB, PHGDH, and PSAT1 in LV was signatory for the BCAA metabolism pathway (Fig 5C). The unbiased search on MeSH disease terms indicated that altered genes are highly associated with HF, cardiomyopathy, hypertrophy, cardiac injury, hyperglycemia, hyperinsulinism, and lipid metabolism errors (Fig 5D). The gene expression profile reflects the crosstalk of obesity, diabetes, and CVD (Fig 5E). For instance, uncoupling protein 3 (UCP3), carnitine palmitoyltransferase 1A (CPT1A), and patatin like phospholipase domain containing 2 (PNPLA2), which are at the crossroad of defects in lipid metabolism, glucose metabolism, and heart injury, were increased in the LV of obese ZSF1 rats. Elevated levels of PDK4 and thioredoxin interacting protein (TXNIP), and decreased expression of the array of genes associated with both heart diseases and compromised glucose metabolism were observed (Fig 5E). Interestingly, altered genes and signaling pathways were also shared by different subtypes of CVD (Fig 5F). For example, decreased MYL2 and MYH6 gene expression coincided with increased RYR2, HCN4, CORIN, NPPA, ERBB2, and MYH7 gene expression in hypertrophic and/or dilated cardiomyopathy (Fig 5F).

**Fig 5.**
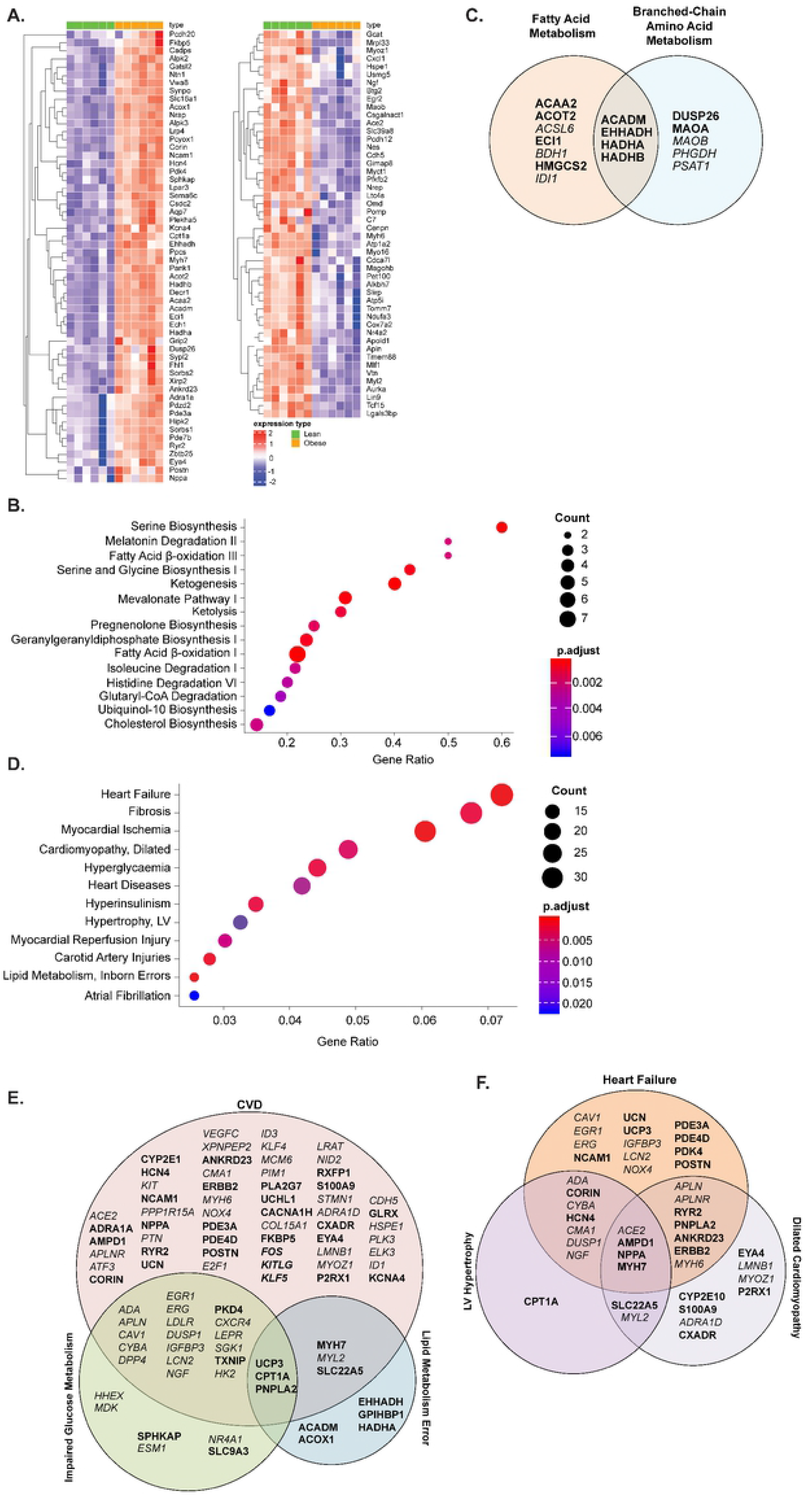
Gene expression analysis of RNA extracted from left ventricle (LV) of 20-week-old obese and lean ZSF1 male rats. The gene expression analysis of RNA revealed increased expression of 56 heart abundant genes and decreased expression of 48 heart abundant genes; the heatmap represents the individual expression changes in obese vs lean ZSF1 rats (A). Ingenuity pathway analysis (IPA) of enriched metabolic pathways (B) and comparison of differentially expressed genes (DEGs) presented in the enriched metabolic pathways from IPA over-represented cardiometabolic Medical Subject Headings (MeSH) terms (C). IPA of 267 significantly increased (FC > 1.5, Benjamini-Hochberg corrected p-value < 0.05) and 431 significantly decreased genes (FC < 2/3, p-value < 0.05) in LV of obese ZSF1 rats (D); comparison of DEGs enriched in cardiovascular disease (CVD), impaired glucose metabolism, and lipid metabolism error disease pathways (E); and comparison of DEGs associated with different subgroups of heart disease pathways from over-representative MeSH term analysis (F). Gene symbols in bold indicate increase of expression and in italic indicate decrease of expression.

Together, these data provide a compelling case that obese ZSF1 rats exhibit multiple features of human CMS, including pathological changes in circulating biomarkers and in the expression of heart abundant genes, cardiovascular dysfunction with preserved ejection fraction, and decreased exercise capacity.

### Obese ZSF1 male rats treated with Fc-GDF15 for 12 weeks demonstrated significant metabolic improvement by changes in systemic parameters and biomarkers of obesity and metabolism impairments

For the duration of Fc-GDF15 or vehicle treatment, both groups of obese ZSF1 rats were monitored weekly for food intake and body weight and biweekly for blood levels of total cholesterol, triglycerides, insulin, and glucose (Fig 6). Within the first 2 weeks of Fc-GDF15 treatment, body weight (Fig 6A), food intake (Fig 6B), blood triglycerides (Fig 6C), and blood glucose levels (Fig 6D) decreased consistently until the end of the study. Blood cholesterol was 20–30% lower in the Fc-GDF15–treated obese ZSF1 rats (vs vehicle-treated animals) from week 1 to week 12 of the treatment; however, this decrease did not achieve significance at any tested time point of the study (Fig 6E). At the same time, insulin levels were significantly increased at weeks 9 and 12 of Fc-GDF15 treatment (Fig 6F). At the end of the 12-week Fc-GDF15 treatment, circulating adiponectin concentration was significantly increased (Fig 6G). Also, endogenous rat GDF15 levels in the serum of Fc-GDF15–treated and vehicle-treated obese ZSF1 rats remained similar (Fig 6H).

**Fig 6.**
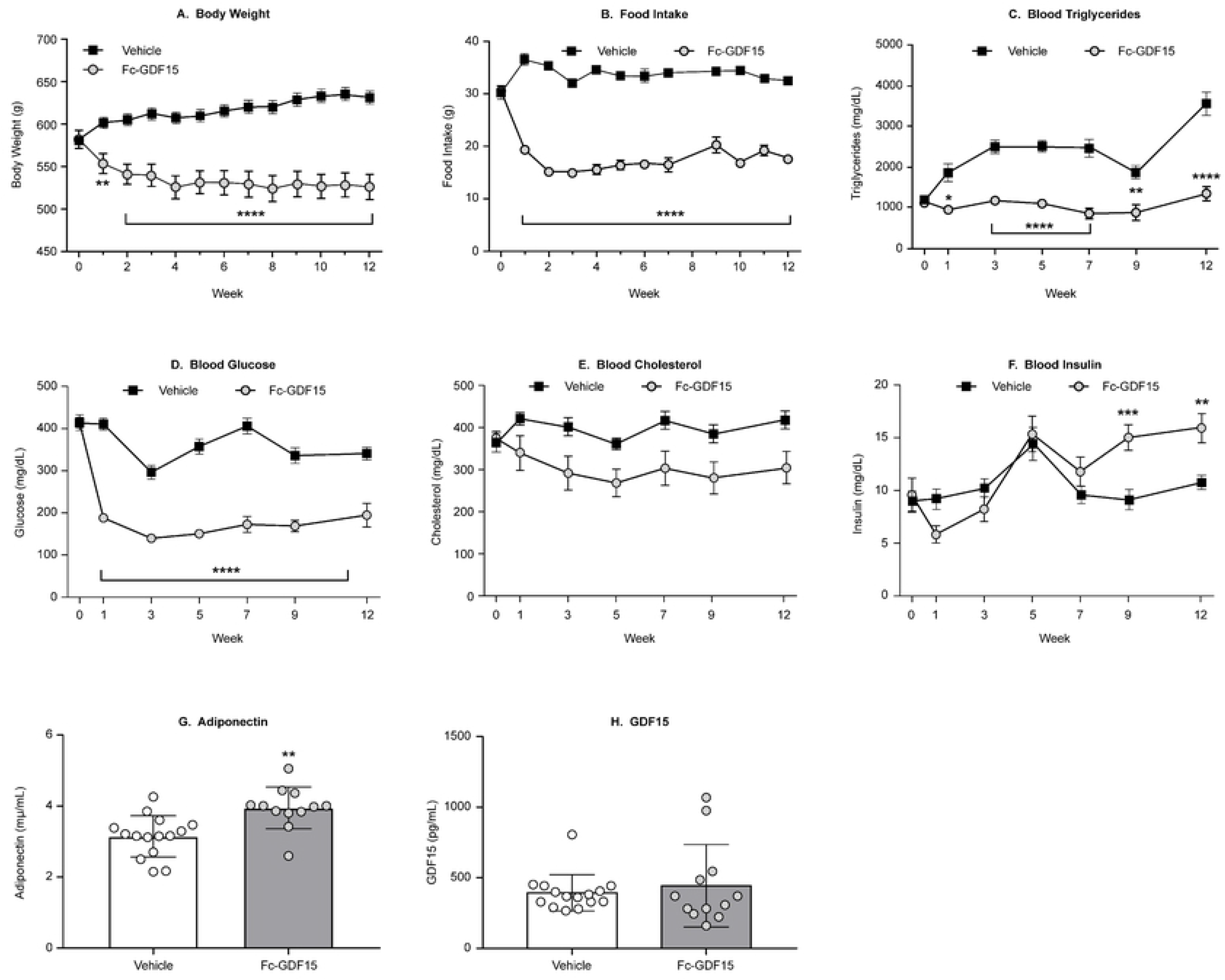
Effect of treatment of ZSF1 obese rats with Fc-GDF15 on systemic metabolic biomarkers. Treatment of ZSF1 rat with Fc-GDF15 resulted in decreased body weight (A) and food intake (B), decrease in blood triglyceride (C) and glucose levels (D), and trend in decrease of blood cholesterol (E). Blood insulin increased over time with GDF15 treatment (F). Twelve weeks of Fc-GDF15 treatment resulted in increased blood adiponectin (G) and had no effect on endogenous levels of rat GDF15 (H). Stars (*) indicate significance (* *p* < 0.05, ** *p* < 0.01, *** *p* < 0.001, **** *p* < 0.0001) by unpaired two-tailed *t* test.

### Fc-GDF15 treatment of obese ZSF1 rats led to significant changes in systemic levels of CVD-related biomarkers

The systemic levels of cardiovascular biomarkers in obese ZSF1 rats were evaluated at the end of the study and are presented in S4 Table. Following 12 weeks of Fc-GDF15 administration, we observed significant changes in the array of cardiovascular circulating biomarkers. Thus, systemic markers of cardiac injury vWF (Fig 7A) and Myl3 (Fig 7B) were 30–50% lower (*p* < 0.05) in the Fc-GDF15 group. HF-associated OPN (Fig 7C), markers of vascular injury sE-selectin (Fig 7D), sICAM (Fig 7E), VEGF (Fig 7F), and marker of cardiovascular fibrosis TIMP-1 (Fig 7G) were significantly lower in the serum/plasma of Fc-GDF15–treated vs vehicle-treated obese ZSF1 rats (*p* < 0.05). At the same time, the other systemic biomarkers of CVD, such as NT-proBNP, NT-proANP, BNP, FABP3, MCP1, and aldosterone were not significantly changed by the 12-week–long Fc-GDF15 treatment (S4 Table).

**Fig 7.**
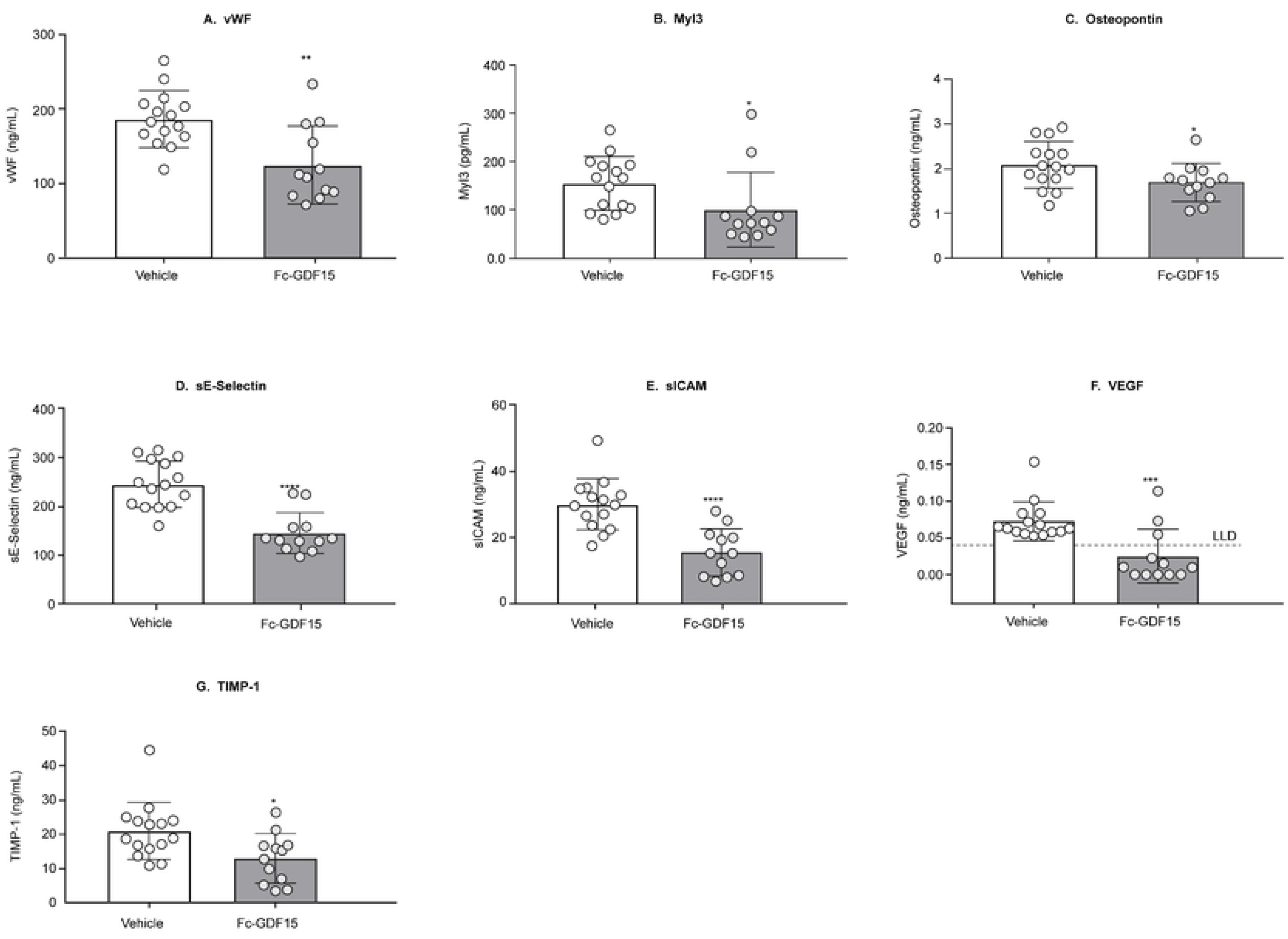
Effect of Fc-GDF15 treatment of ZSF1 obese rats on systemic CVD biomarkers. Obese ZSF1 rats treated with Fc-GDF15 for 12 weeks exhibit decrease in systemic levels of cardiovascular disease–related biomarkers vWF (A), Myl3 (B), osteopontin (C), sE-selectin (D), sICAM (E), VEGF (F), and TIMP-1 (G) at the end of the study. Stars (*) indicate significance (* *p* < 0.05, ** *p* < 0.01, *** *p* < 0.001, **** *p* < 0.0001) by unpaired two-tailed *t* test. *N* = 11 for vehicle-treated group and *N* = 8 for Fc-GDF15–treated group.

### Fc-GDF15 therapeutic approach affected echocardiography parameters of left heart function and improved exercise capacity in obese ZSF1 rats

When we characterized obese ZSF1 rats, we demonstrated that at the age of 20 weeks, males exhibited impaired cardiac function and decreased exercise capacity. Twelve weeks of Fc-GDF15 treatment of obese ZSF1 male rats resulted in significantly decreased heart-to-brain weight ratio (by echocardiography; S5 Table, Fig 8), LV mass, cardiac output, stroke volume, and ejection fraction when compared with obese ZSF1 rats treated with vehicle (Fig 8A–8E). Fc-GDF15 treatment had no effect on heart rate (Fig 8F). After 12 weeks of treatment, when exposed to a treadmill challenge, the Fc-GDF15 group demonstrated improved exercise capacity (S5 Table) by exhibiting significantly longer running time (Fig 8G) and running distance (Fig 8H) vs the vehicle group. However, peak VO_2_ (a measure of aerobic fitness) and RER (a fatigue measure) were not significantly different between groups (S5 Table), suggesting that both groups were run to the same level of exhaustion.

**Fig 8.**
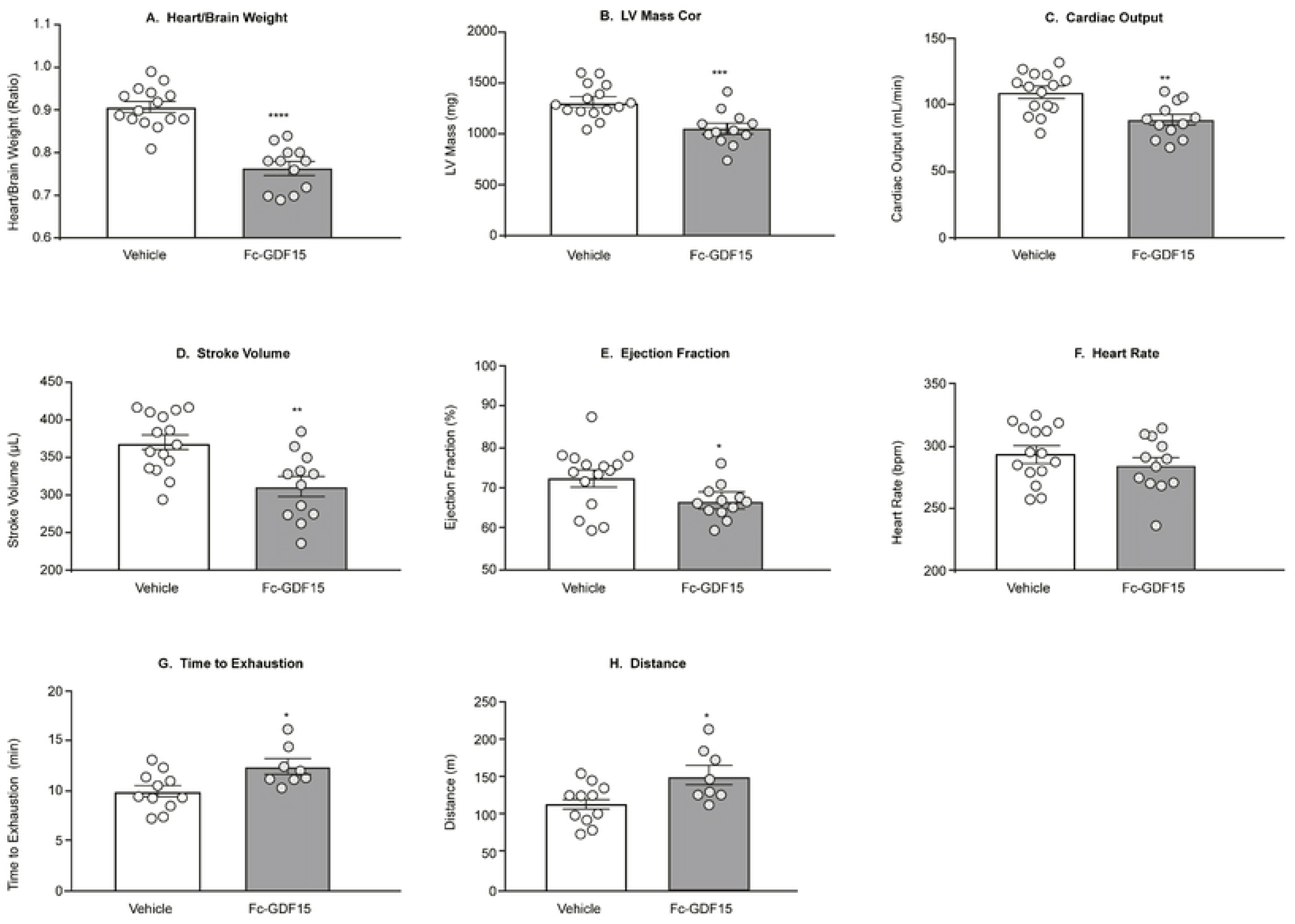
Fc-GDF15–treated obese ZSF1 rats demonstrated improved parameters of cardiac function and increased exercise capacity. Twelve weeks of Fc-GDF15 treatment decreased heart weight (A); decreased left ventricular (LV) mass cor (B), cardiac output (C), stroke volume (D), and ejection fraction (E) but not heart rate (F). Fc-GDF15–treated obese ZSF1 rats demonstrated improved exercise capacity during treadmill exercise by increased length of time to exhaustion (G) and running distance (H). Stars (*) indicate significance (* *p* < 0.05, ** *p* < 0.01, *** *p* < 0.001, **** *p* < 0.0001) by unpaired two-tailed *t* test.

## Discussion

CMS is a combination of primarily IR-associated metabolic disorders with a reciprocal relationship between impaired metabolism and HFpEF [3]. Our understanding of the pathophysiology and mechanisms of CMS-related HFpEF is limited due to the minimal availability of human myocardial biopsies and the lack of animal models that closely mimic human pathology.

Recently, obese ZSF1 rats were proposed as a robust CMS model because at the age of 20 weeks, males demonstrate hypertension, obesity, T2D, impaired metabolism, HF, and exercise intolerance [4, 7, 22, 30]. The LV hypertrophy, LA dilation, and increased myocardial stiffness due to myofilament changes (without significant interstitial fibrosis) are well documented in obese ZSF1 rats [8]. In our study, we have confirmed the previous observations (Figs 1 and 3). Additionally, we have examined systemic levels of pancreas-related biomarkers in obese ZSF1 rats and demonstrated pancreatic beta-cell dysfunction with significantly increased serum C-peptide, proinsulin, glucagon, and amylin (Fig 2A–2D). Since renal dysfunction is a part of CMS, Hamdani *et al*. [8] compared major parameters of kidney function in 20-week-old obese vs lean ZSF1 rats and reported hyperglycemia caused glycosuria, increased urine output, compensatory water intake, and proteinuria, suggesting the presence of diabetic nephropathy despite preserved creatinine clearance and plasma protein levels. Significantly increased systemic kidney injury biomarkers KIM-1, NGAL, and clusterin (Fig 2E–2G) confirmed the presence of diabetic nephropathy and kidney damage in 20-week-old obese ZSF1 male rats.

Since biomarkers have emerged as powerful diagnostic and/or prognostic tools for the variety of CVD in humans, we studied and analyzed in depth the milieu of systemic protein and local molecular cardiovascular biomarker changes (LV of the heart) triggered by the development of CMS in obese ZSF1 rats and evaluated the translational value of the observations. Notably, among the array of systemic changes we have documented is an increase in aldosterone (Fig 4A) together with significant attenuation of NT-proBNP (Fig 4E). The reciprocal relationship between systemic aldosterone and NT-proBNP was recently reported in a human CMS populational study (*n* = 1674; age ≥ 45 years) [31], where the authors demonstrated strong (*p* < 0.001) association of aldosterone increase with hypertension, obesity, chronic kidney disease, metabolic syndrome, high triglycerides, concentric LV hypertrophy, and atrial fibrillation. They also demonstrated reverse correlation between aldosterone and NT-proBNP. During the last decade, several scientific reports from human populational studies have demonstrated that serum NT-proBNP concentrations were relatively lower in overweight and obese patients [32–34], suggesting that natriuretic peptides may provide a link between the heart and adipose tissue [35]. The observation that human subcutaneous adipose tissue from obese subjects with T2D exhibits markedly increased natriuretic peptide (NP) receptor expression and higher clearance of NT-proBNP from the circulation [36] may explain why we observed a significant decrease in serum NT-proBNP without NPPB mRNA expression changes in the left heart of obese ZSF1 rats (Fig 4).

FABP3 is abundantly expressed in cardiomyocyte cytoplasm; its circulating level is positively correlated with LV mass index (*p* < 0.0001, *r* = 0.7226). Therefore, FABP3 was proposed as an early biomarker of myocardial injury in humans [37]. ST2, a circulating protein marker of cardiomyocyte stress and fibrosis, which increases in patients across a wide spectrum of CVDs, is now recommended by the American College of Cardiology Foundation and American Heart Association joint guidelines for additive risk stratification in patients with HF [38]. Chronic inflammation contributes to cardiac fibrosis, and systemic IL-16 levels are specifically elevated in HFpEF patients (compared with HF with reduced ejection fraction [HFrEF] and controls), with a significant association between serum IL-16 and indices of LV diastolic dysfunction (LAVI, E/E’, and DWS) [39]. In concert with the published results from human studies, serum FABP3, ST2, and IL-16 protein concentrations were significantly elevated in 20-week-old obese ZSF1 male rats (Fig 4B–4D).

OPN (abundant kidney protein majorly synthesized in the loop of Henle) is among the biomarkers of HF progression; it is known to be expressed at medium levels in heart tissue by endothelial cells, cardiomyocytes, and fibroblasts. Several groups reported human plasma OPN elevation in patients with advanced HF [40] and specifically in HFpEF cohorts [41]. Lopez *et al*. [42] have reported no association of plasma OPN with HF, whereas the myocardial expression of OPN was highly elevated in HF patients (*p* < 0.0001). Our preclinical results confirm the latter report and provide, for the first time, evidence that myocardial OPN is up-regulated in the heart of rats with HFpEF, as shown in murine models of HF [43].

Based on transcriptome analysis, approximately 60% (*n* = 12,224 of 19,613) of protein-coding genes are expressed in heart tissue [44]. Two hundred of these genes show an elevated expression in heart compared to other tissue types, and most of the corresponding proteins (localized in the cytoplasm and in the sarcomeres) are involved in muscle contraction, ion transport, and ATPase activity. Understanding the pathophysiology of cardiac alterations in CMS is critical. Our comparative RNA-seq analysis (lean vs obese ZSF1) of rat LV heart abundant genes revealed 267 significantly increased genes and 431 significantly decreased genes, of which 56 increased genes and 48 decreased genes are among the abundant genes in human heart (Fig 5A). Enriched signaling and disease pathway analyses added to our current understanding of how intermediary metabolism affects LV hypertrophy and influences heart tissue remodeling and repair. In the ZSF1 rat CMS model, obesity and impaired metabolism have also greatly increased CVD pathology by altering two major metabolic pathways in heart tissue (Fig 5B and 5C)—fatty acid metabolism and BCAA metabolism. In general, cardiac metabolic homeostasis of fatty acid, glucose, ketone bodies, and BCAAs is well established through intertwined regulatory networks [45, 46]. Thus, gene expression of essential enzymes involved in both fatty acid metabolism and BCAA catabolism—ACADM, EHHADH, HADHA, and HADHB—is greatly increased in obese ZSF1 LV heart tissue. It has been reported that acyl-CoA deficiency in human failing heart disrupts cardiac energy production and leads to cardiac lipotoxicity, which has a negative impact on the heart as it impairs its ability to function and pump properly [47]. ACAA2, ACADM, ACSL6, ACOT2, EHHADH, HADHA, and HADHB are enzymes functionally involved in acyl-CoA metabolism. Human genome-wide association studies (GWAS) have indicated that mutations in HADHA and HADHB are associated with familial hypertrophic cardiomyopathy [48], and mutations in CPT1, ACADM, and ACAA2 genes are associated with impaired mitochondrial fatty acid β-oxidation [49]. Several enzymes (ACADM, ACSL6, EHHADH, and HMGCS2) are regulated by proliferator-activated receptors (PPARs), which manipulate the fuel supply and substrate and modulate HF progression [50, 51]. Based on the MeSH database analysis, we have demonstrated that in the obese ZSF1 rat model, the top representative diseases were CVD, hyperinsulinemia, hyperglycemia, and lipid metabolism defects (Fig 5D). Obesity and T2D are among the top drivers of HFpEF progression [2, 3]. Transcriptome profiling revealed the array of genes and pathways potentially mediating the cardiac metabolic shift, glucose and lipid toxicity, and the development of cardiomyopathy (Fig 5E). For instance, UCP3, CPT1A, and PNPLA2 are shared DEGs among impaired lipid metabolism, glucose metabolism, and CVD. CPT1A and UCP3 are involved in cardiac glucose oxidation, mitochondrial fatty acid oxidation, and ATP production. CPT1 inhibition has demonstrated beneficial effect in HF [52], and UCP3 was considered a marker for cellular metabolic state [53]. Mutations in PNPLA2 (which encodes adipose triglyceride lipase, ATGL) have been associated with triglyceride deposit cardiomyovasculopathy (TCGV) [54]. Mori *et al*. [55] have found that deletion of pyruvate dehydrogenase kinase 4 (PDK4) prevented angiotensin II-induced cardiac hypertrophy and improved cardiac glucose oxidation and energy usage. MeSH-based disease analysis demonstrated that significant changes in cardiomyocyte genes (decreased MYL2 and MYH6 and increased RYR2, HCN4, CORIN, NPPA, ERBB2, and MYH7) in LV of obese ZSF1 rats were strongly associated with dilated cardiomyopathy, LV hypertrophy, and HF (Fig 5F), and were similar to those observed in human genetic studies [56]. The array of expression changes in LV heart abundant genes in obese ZSF1 rats would help us to better understand the cardiac metabolic shift and cardiomyopathy in CMS; to allocate the major players mediating the crosstalk between glucose, lipid metabolism, and development of CVD; and to potentially identify key markers of cardiac energy status and heart injury grade. Hence, transcriptome profiling of lean and obese ZSF1 rats further supports the translational value of the obese ZSF1 CMS rat model in biomarker discovery and evaluation of therapeutic targets.

GDF15 is a distant member of the TGF-β superfamily. It is secreted, circulating in plasma as a 25 kDa homodimer [9, 10], and has become a novel exploratory biomarker of CMS because its circulating levels are increased in humans with metabolic syndrome [11, 14, 19] and in subjects with increased risk of CVD [12, 13]. In concert with human data, GDF15 levels do rise following a sustained high-fat diet or dietary amino acid imbalance in mice [57], and according to our present report (Fig 1G), GDF15 is significantly increased in the serum of obese ZSF1 rats. To date, GDF15 is positioned as a stress-induced hormone that mediates an aversive dietary response in preclinical species; when it was tested in obese mouse and non-human primate models, treatment resulted in significantly reduced body weight and food intake, increased energy expenditure, and improved glucose tolerance [14, 15].

Results reported here make a compelling case for obese ZSF1 rats exhibiting multiple features of human CMS, which include pathological changes in systemic renal, metabolic, and CVD circulating biomarkers, HFpEF, and decreased exercise capacity. Recently published studies in obese preclinical models [14, 15, 57] demonstrated aversive dietary response to GDF15 treatment, leading to improvement in metabolic parameters. We treated obese ZSF1 rats with Fc-GDF15 for 12 weeks and compared the outcome with that in vehicle-treated obese ZSF1 rats. Twelve-week Fc-GDF15 treatment resulted in a significant decrease in body weight and food intake, demonstrating its capacity to mediate aversive dietary response. Fc-GDF15 improved metabolic parameters (decreased blood glucose and triglycerides) in obese ZSF1 rats (Fig 6A–6F), similar to previously reported results in diet-induced obese mice and obese cynomolgus monkeys [15], and significantly increased serum adiponectin in Fc-GDF15– treated obese ZSF1 rats. Adiponectin is mainly secreted by adipocytes, but also by skeletal muscle cells, cardiac myocytes, and endothelial cells. Reduction of adiponectin plays a central role in CMS because it is positively associated with insulin sensitivity [58] and shows anti-atherogenic and anti-inflammatory properties; according to numerous epidemiological studies, hypoadiponectinemia (adiponectin deficiency) is additionally associated with CVDs such as hypertension, coronary artery disease, and LV hypertrophy [59]. While circulating adiponectin was decreased by 30% in obese vs lean ZSF1 rats (Fig 1F), Fc-GDF15 treatment led to its significant systemic increase (29%; Fig 6G) vs obese ZSF1 rats treated with vehicle. Although Fc-GDF15 treatment did not lead to systemic changes in aldosterone, NT-proBNP, and FABP3 (S4 Table) in obese ZSF1 rats, the novel and exploratory systemic protein markers [60] of cardiac injury vWF and Myl3; HF-associated OPN; markers of vascular injury sE-selectin, sICAM, and VEGF; and cardiovascular fibrosis marker TIMP-1 were significantly lower in Fc-GDF15–treated obese rats vs vehicle-treated obese rats (Fig 7). In addition to the positive changes in the array of systemic cardiometabolic biomarkers, 12-week–long Fc-GDF15 therapy led to a considerable decrease in heart weight, improved parameters of LV function (decreased LV mass, cardiac output, and stroke volume; Fig 8), and increased exercise capacity (time to exhaustion and distance; Fig 8) when compared with vehicle-treated obese ZSF1 rats.

## Conclusion

In summary, the obese ZSF1 male rat represents a preclinical model with established HFpEF that can mimic human CMS. Furthermore, Fc-GDF15 treatment of obese ZSF1 rats demonstrated its cardioprotective therapeutic effect in this model. These findings may have important clinical implications for potential pharmacologic treatment of obesity and associated comorbidities such as HF, where the unmet medical need remains high.

## Acknowledgements

The authors thank Cathryn M. Carter of Amgen Inc for editorial support; she received compensation as an employee of Amgen Inc.

## Author Contributions

Marina Stolina designed the studies, analyzed the results and wrote the manuscript; Xin Luo provided transcriptome analysis of RNAseq results, differential expression pathway analysis and contributed to the Results and Discussion of current manuscript; Denise Dwyer produced, and analyzed biochemical biomarkers–related data; Chun-Ya Han characterized ZSF1 strain based on the local and systemic biomarker profiles; Rhonda Chen and Ying Zhang provided echocardiography data collection and initial analysis; YuMei Xiong designed and provided initial analysis of Fc-GDF15 study; Yinhong Chen provided echocardiography and PV loop data collection and comprehensive analysis; Jun Yin provided transcriptome analysis and initial interpretation of RNAseq results; Brandon Ason provided invasive hemodynamic assessment, treadmill assessment and interpretation of the studies; Clarence Hale contributed to the studies design, data analysis and interpretation and the writing of the manuscript; Murielle M. Véniant provided major input to the studies design, analysis and interpretation of the results and the writing/editing of the manuscript.

All authors read and approved the final manuscript.

## Supporting information

**S1 Text. Gene models used for alignment and quantification.**

**S1 Table. Metabolic and renal biomarkers in serum/plasma of 20-week-old lean and obese ZSF1 male rats.**

**S2 Table. Cardiovascular biomarkers in circulation of 20-week-old lean and obese ZSF1 male rats.**

**S3 Table. Invasive hemodynamic assessment, echocardiography, and exercise capacity of 20-to 21-week-old lean and obese ZSF1 male rats.**

**S4 Table. Effect of 12-week–long Fc-hGDF15 treatment on systemic levels of cardiovascular markers.**

**S5 Table. Effect of 12-week–long Fc-hGDF15 treatment on parameters of invasive hemodynamic assessment, echocardiography, and exercise capacity of obese ZSF1 male rats.**

**S6 Table. Heart abundant tissue gene expression increase in LV of obese vs lean ZSF1 groups.**

**S7 Table. Heart abundant tissue gene expression decrease in LV of obese vs lean ZSF1 groups.**

